# Evaluation of the Safety and Efficacy of the Respiratory Syncytial Virus FG Chimeric Vaccine KD-409 in Rodent Models for Maternal and Pediatric Vaccination

**DOI:** 10.1101/2025.02.12.637802

**Authors:** Ryo Yamaue, Madoka Terashima, Kenji Soejima, Masaharu Torikai

**Affiliations:** KM Biologics Co., Ltd., Kikuchi Research Center, 1314-1 Kyokushi kawabe, Kikuchi-shi, Kumamoto 869-1298, Japan

## Abstract

Respiratory syncytial virus (RSV) causes severe infection in neonates and infants. However, a suitable RSV vaccine for children has yet to be approved. We developed an effective and safe RSV vaccine for newborns and children. Accordingly, we evaluated the safety and efficacy of the RSV FG chimeric protein KD-409, which incorporates a highly conserved region of the RSV G protein into the RSV F protein, in several rodent models. We confirmed the effect of our vaccine-induced antibody transfer using a guinea pig model. Subsequently, we evaluated the exacerbation of infection in a BALB/c mouse model of passive immunity, which was designed to mimic the vaccination of pregnant women. Notably, KD-409 in the presence of alum did not exacerbate infection, which occurred upon administering pre-F with alum. Our active immunization model of BALB/c mice, which simulated vaccination with a pediatric vaccine, suggested that KD-409 with alum was less likely to exacerbate inflammation than FI-RSV or pre-F with alum. The efficacy was evaluated in a cotton rat model, in which KD-409 demonstrated better protection against infection than pre-F without adjuvant, the only currently approved formulation for immunizing pregnant women. Collectively, antibody transfer in pregnant guinea pigs during immunization, safety in mice during passive and active immunization, and efficacy in cotton rats demonstrate the high potential of KD-409 as a novel vaccine candidate. Our results provide insights into the potential of KD-409 as a safe and effective next-generation RSV vaccine that can cover the neonatal-to-pediatric age range.

**IMPORTANCE:** The safety and efficacy of KD-409, an RSV FG chimeric protein, were evaluated in appropriate rodents. We found that KD-409 was less likely to exacerbate infections and symptoms, and exhibited superior safety and efficacy to pre-F without adjuvant, which is the only approved vaccine for pregnant women. Thus, KD-409 represents a valuable immunization modality for maternal and pediatric immunization.

## INTRODUCTION

Respiratory syncytial virus (RSV) infection is more severe in neonates and infants. In 2019, an estimated 3.6 million RSV-acute lower respiratory infection (ALRI) hospitalizations and 101,400 RSV-ALRI deaths occurred in children aged 0–60 months worldwide (1), with an estimated 1.4 million RSV-ALRI hospitalizations and 45,700 RSV-ALRI-attributable deaths reported in infants aged 0–6 months (1). RSV vaccines for children were developed in the 1960s using FI-RSV as an antigen; however, the vaccinated group exhibited adverse events such as vaccine-enhanced disease (VED) (2). Despite the challenges associated with RSV vaccine development for half a century, the strategy of preventing infection with transferable antibodies after vaccination of pregnant women has been successful. The Food and Drug Administration approved the first RSV vaccine for neonates on August 21, 2023 (3). Nevertheless, a phase III clinical trial (NCT04424316) in pregnant women and neonates failed to meet all primary endpoints, leaving room for improvement in efficacy (4, 5). Additionally, RSV vaccine development for children is ongoing, with clinical trials focusing on live attenuated vaccine candidates. However, suitable candidates are yet to be selected.

In the development of RSV vaccines, safety assurance is essential due to concerns regarding VED. Three causes of symptom exacerbation in natural infection after FI-RSV vaccination have been reported: i) low avidity of induced antibodies (6); ii) Th2 induction by carbonylated proteins resulting from formalin treatment (7); and iii) Th2 induction by G proteins (8). Issues i and ii are deemed avoidable with the appropriate use of adjuvants, recombinant proteins, or attenuated live viruses. In the case of iii), using only the F protein as an antigen may be a suitable solution, although VED caused by adjuvanted F has been documented in some animal experiments (9). Given that the neutralizing capacity of anti-G protein antibodies is comparable to that of anti-F antibodies (10), G proteins, as immune antigens, appear to be promising in terms of efficacy. The CX3C motif, which functions as both a host cell adhesive and aggravating factor, is located at residues G182–186 within the conserved central domain (CCD; G157–198), surrounded by cysteine loops (G173–186) and a highly conserved region (11, 12). A clinical trial of BBG2Na, which fused the RSV A G130–230 residues (G2Na) containing this CCD region to the albumin-binding domain of streptococcal G protein (BB), was terminated following the occurrence of unexpected side effects (13).

Although the pre-F protein antigen vaccine is promising, several studies have indicated increased inflammation following RSV exposure after immunization (9). Understanding the inflammatory response is crucial for developing RSV vaccines. Severe cases of neonatal RSV infection are common up to 6 months of age, with a peak at around 2–3 months of age (14–16). The half-lives of antibodies induced by vaccination of pregnant women with diphtheria, tetanus, and pertussis (DTP) vaccines are 28.7 days (tetanus toxoid antibodies) and 35.1 days (pertactin antibodies) (17), while those of RSV vaccines may be similar. Vaccination of pregnant women alone cannot cover the entire high-risk period of severe disease. Additionally, RSV vaccines may be capable of inducing antibody-dependent enhancement (ADE) of infection, as observed with dengue vaccines, if the antibody concentration in the blood decreases; therefore, this possibility needs to be examined.

We hypothesized that the efficacy and safety of the vaccine could be enhanced by inducing antibodies near the CCD of the G protein rather than just the F protein. KD-409 was designed to induce antibodies around the CX3C motif, as it lacks the CX3C motif and is therefore less likely to cause inflammation. Although not directly binding to the CX3C motif, this anti-G antibody is expected to act as a form of steric hindrance due to the bulkiness of the antibody itself, inhibiting the CX3C-CX3CR1 interaction. In a previous report, we demonstrated that KD-409 outperformed pre-F in protecting against infection in a pregnant mouse model (18). Although our study demonstrated efficacy in a mouse pregnancy model, we were unable to report safety or efficacy data in animals other than mice.

In the present study, we evaluated the safety and efficacy of KD-409 in multiple rodent species. As a novel vaccine encompassing all age groups, i.e., from newborns to children, KD-409 could be a next-generation vaccine with superior safety and efficacy compared to RSV F-based vaccines.

## RESULTS

### Antibody transfer in a guinea pig pregnancy model

The guinea pig is an appropriate model for mimicking human antibody transfer (19). Nonclinical studies on RSV vaccines have been conducted in guinea pigs, and antibody transfer has been confirmed for F protein-only antigens (20).

We employed a guinea pig pregnancy model system to confirm the antibody transfer of KD-409, which is a chimeric FG protein containing a part of the G protein (Fig. 1A). Anti-F antibodies were detected in the serum of guinea pigs that had been immunized twice, as well as in the serum of their pups born to respective mothers on day 0 (day of birth = day 0) (Fig. 1B, E). On day 0, the titers of both anti-F and neutralizing antibodies were higher in the pups than in the mother, indicating that the antibody concentration could be attributed to the transfer of antibodies from the mother to the pups (Fig. 1B, E, H). The dose-dependent induction of anti-F and neutralizing antibodies was confirmed by comparing immunizations with 20 and 5 µg (Fig. 1B–H). These results indicated the occurrence of antibody transfer in the guinea pig model. In humans, antibody transfer to infants occurs through maternal immunity.

**Fig. 1.**
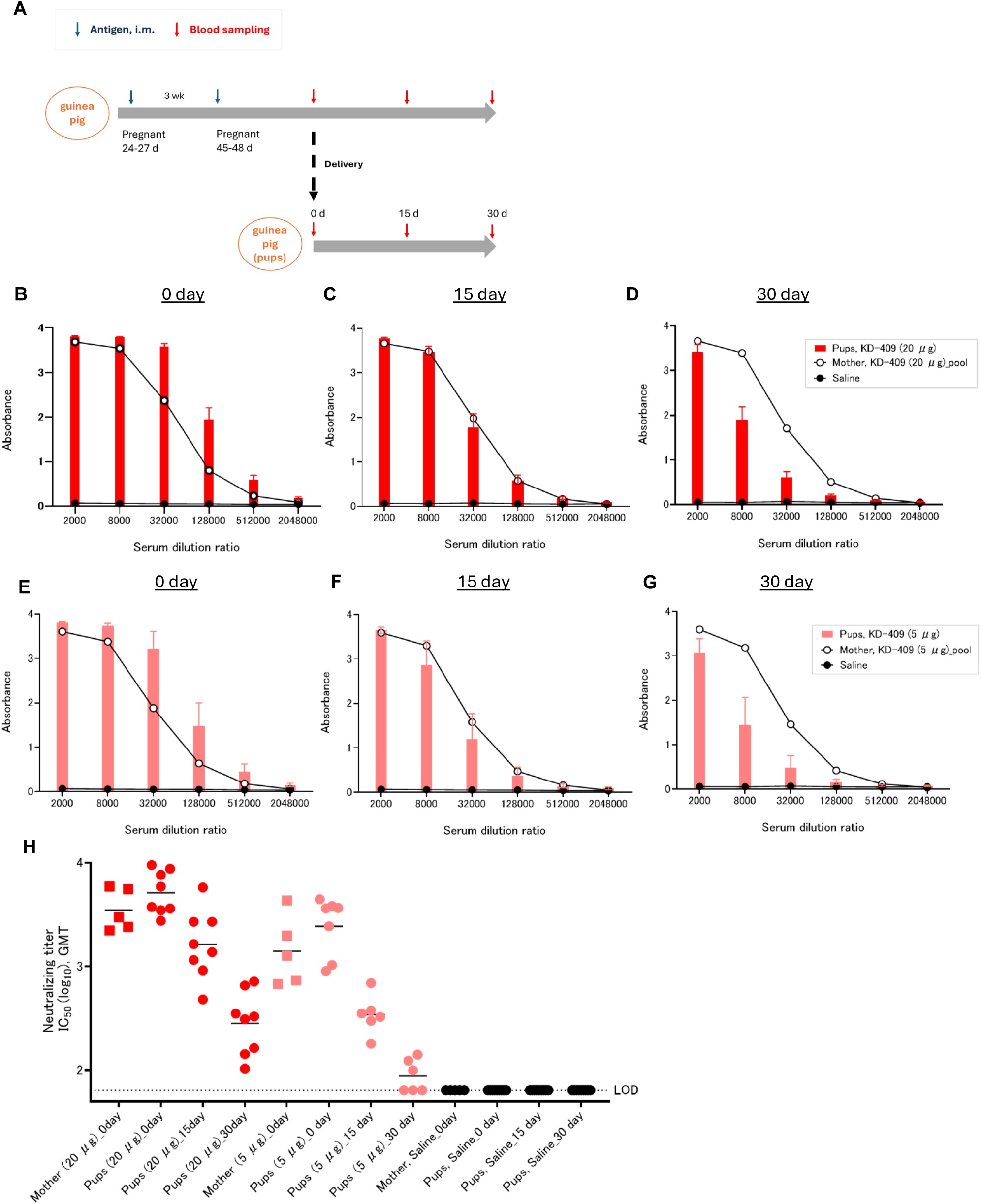
Neutralizing anti-F antibodies transfer from mother to pup in a guinea pig pregnancy model (A) Experimental flow diagram of the guinea pig pregnancy model. Mother and pup sera were collected 3 weeks after two intramuscular immunizations with KD-409 at 3-week intervals. (B–D) Anti-RSV F antibody titer in the pup serum and mothers’ pooled serum (n = 5–9; days 0, 15, and 30) from animals immunized with a 20 µg/dose. (E–G) Anti-RSV F antibody titer in the pup serum and mothers’ pooled serum (n = 5–9; days 0, 15, and 30) from animals immunized with a 5 µg/dose. (H) Neutralizing antibody titer in the pup and mothers’ serum (n = 5–9; days 0, 15, and 30).

### Evaluation of infection exacerbation following passive immunization in mice

In vaccine safety evaluations, it is crucial to examine the potential for adverse events due to VED and ADE. Although the safety of RSV vaccines has been assessed using *ex vivo* infection enhancement validation (21), few studies have evaluated infection enhancement using animal models.

We evaluated the safety of a passive mouse immunization model, assuming the vaccination of pregnant women (Fig. 2A). Considering the immune serum-administered infection model, we obtained serum samples after two immunizations and determined the serum dilution factor at which the neutralizing capacity was reduced. Mice treated with serum diluted to factor 10^8^ were challenged with RSV, and the number of viral copies in the lung tissue was compared. The results showed that pre-F significantly increased (p = 0.0378) the number of viral copies compared to saline, whereas KD-409 showed results comparable to those of saline (Fig. 2B). Overall, this result suggested that pre-F, but not KD-409, exacerbated infection in the mouse passive immunization model.

**Fig. 2.**
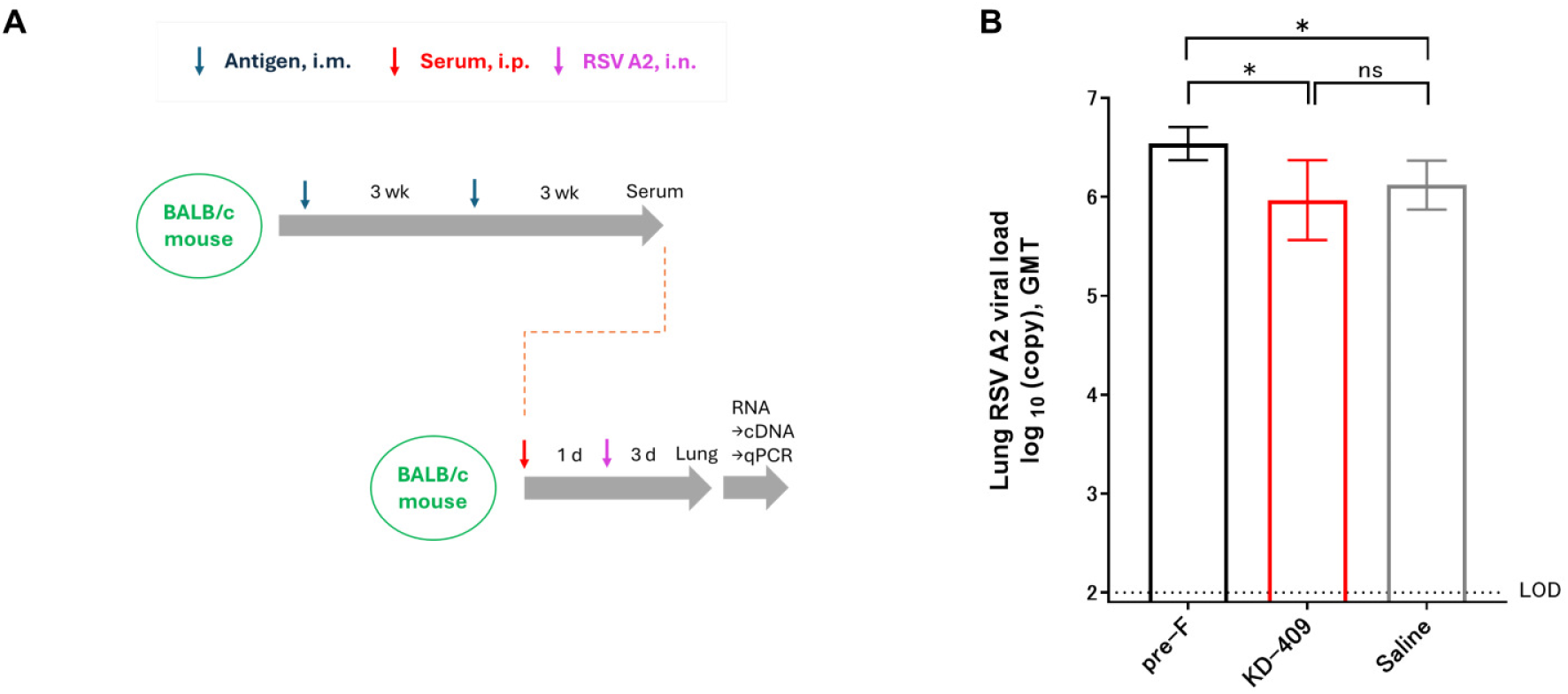
KD-409 with Adju-Phos does not exacerbate infection in passively immunized mice (A) Experimental flow diagram of mouse passive immunization (immunized serum-challenged model). The serum was obtained after two intramuscular immunizations with KD-409 at 3-week intervals. Mice were challenged with RSV 1 day after intraperitoneal administration (i.p.), and lung tissues were harvested 3 days later. (B) Comparison of virus copy numbers of passively immunized mice (immunized serum administration). Statistical analysis was performed by one-way-ANOVA and Dunn’s multiple comparison test. n = 13; *p < 0.05, ns: p> 0.05, not significant. LOD: Limit of detection.

### Evaluation of symptom exacerbation following active immunization in mice

RSV infection in the absence of adequate immunity or weakened blood antibody titers may result in the exacerbation of infection and symptoms. We performed a safety evaluation using an active immunization model in which BALB/c mice were immunized with a low dose of RSV (Fig. 3A). We analyzed the number of viral copies and found that FI-RSV immunization resulted in infection exacerbation (Fig. S1). Given that infection was exacerbated upon administering a 50 ng/dose of FI-RSV containing Adju-Phos, the antigen protein dose was set at 0.5, 5, 50, or 500 ng. Upon examining lung tissue section images, we found that FI-RSV and pre-F showed signs of lung tissue inflammation, whereas KD-409 did not induce any inflammation under these conditions (Fig. 3B). Additionally, the number of polymorphonuclear (PMN) leukocytes, eosinophils, and neutrophils in the bronchoalveolar lavage fluid (BALF) was significantly higher in the pre-F with Adju-Phos group than the saline groups; however, those of KD-409 with Adju-Phos were similar to those in the saline group (Fig. 3C, 3D, 3E). These results suggested that pre-F with Adju-Phos is more likely to exacerbate symptoms in the absence of adequate immunity, whereas KD-409 is less likely to exacerbate symptoms.

**Fig. 3.**
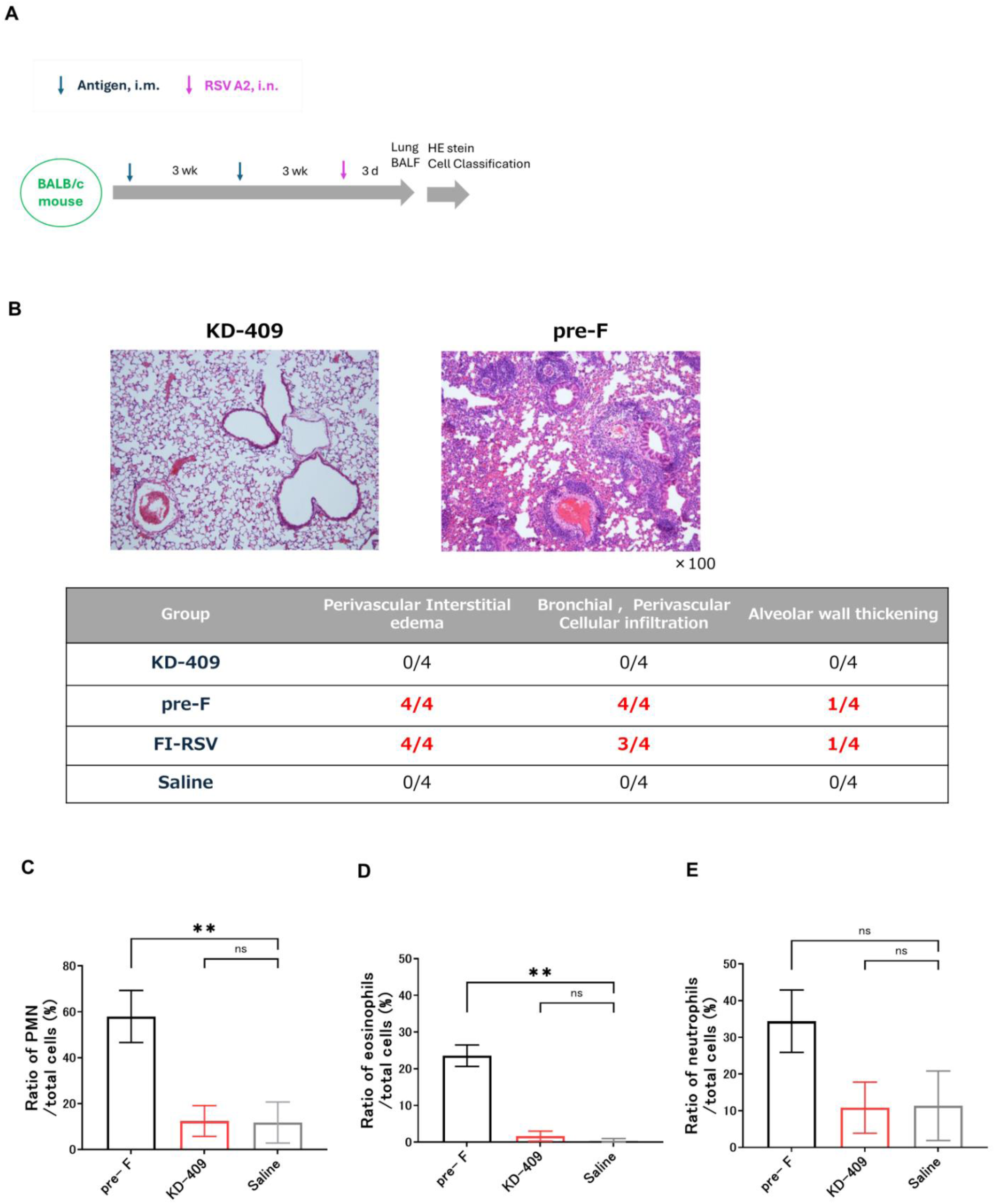
KD-409 with Adju-Phos does not exacerbate symptoms in actively immunized mice (A) Experimental flow diagram of mouse active immunization. After two immunizations with KD-409 at 3-week intervals, mice were challenged with RSV, and the lungs were harvested 3 days later. (B) Representative images of hematoxylin-eosin-stained tissue sections and evaluation of inflammation in lung tissue sections. The proportion of mice showing the most symptoms of exacerbated inflammation (n = 4; 0.5, 5, 50, and 500 ng/dose). (C–E) Ratio of polymorphonuclear (PMN) leukocytes, eosinophils, and neutrophils in the bronchoalveolar lavage fluid (BALF) (n = 3). Statistical analysis was performed by one-way-ANOVA and Dunnett T3 multiple comparison test. **p < 0.01, ns: p > 0.05, not significant.

### Efficacy evaluation by active immunization of cotton rats

We previously evaluated protection against infection using a mouse model (18). Herein, we evaluated efficacy in a cotton rat model, which allows a higher degree of extrapolation to humans (Fig. 4A). Upon incorporating the Adju-Phos adjuvant in the formulation, we found that pre-F and KD-409 were comparable in terms of efficacy (data not shown); however, the vaccine approved for pregnant women was a formulation comprising pre-F without adjuvant. KD-409 with Adju-Phos had a low risk of infection and symptom exacerbation (Figs. 2 and 3). While the safety of KD-409 has been confirmed, the efficacy of KD-409 has yet to be compared to that of adjuvant-free pre-F.

**Fig. 4.**
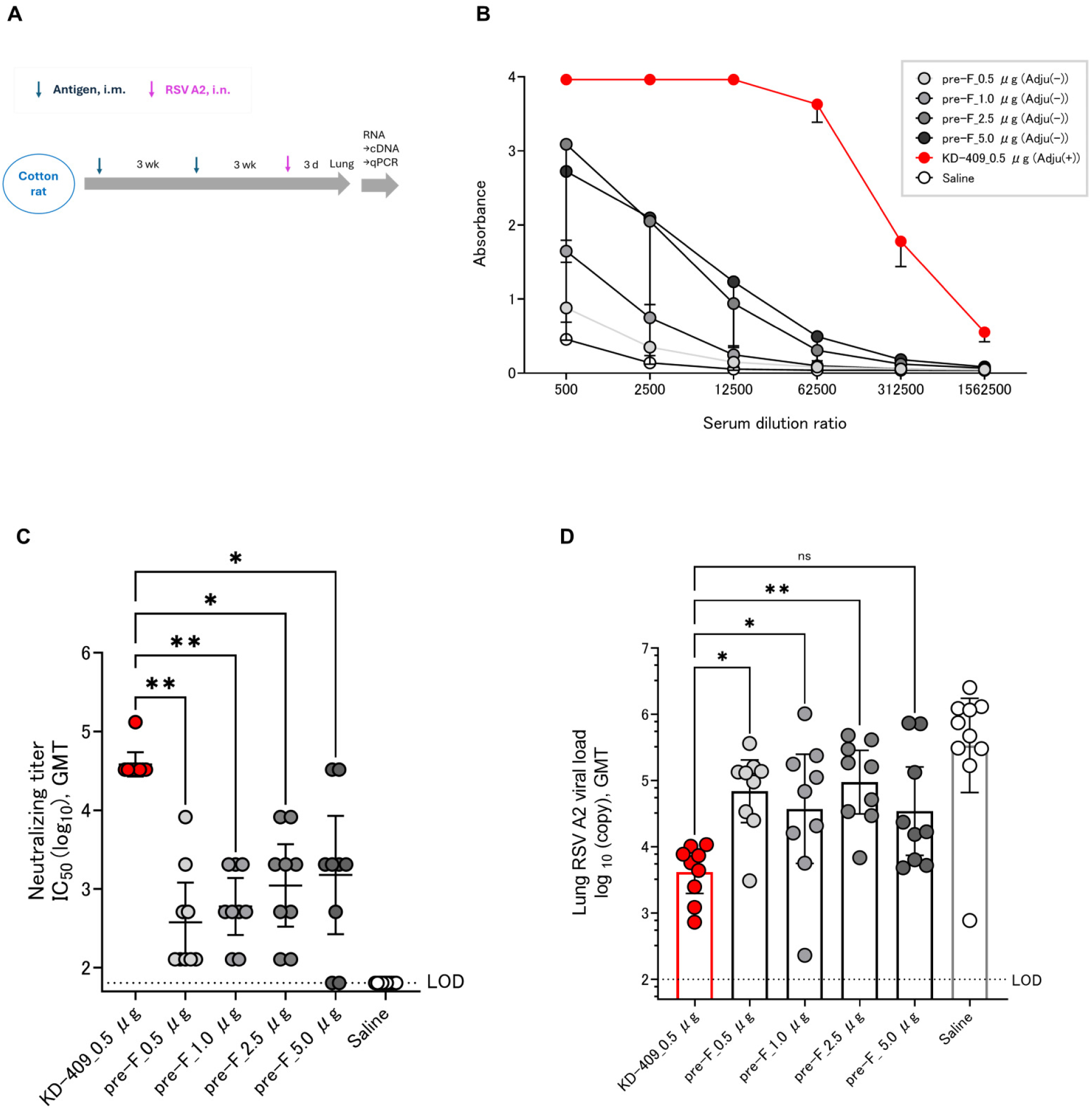
Low-dose KD-409 with Adju-Phos has greater efficacy than adjuvant-free high-dose pre-F in the cotton rat model (A) Experimental flow diagram of the cotton rat model; following two intramuscular injections of KD-409 at 3 week-intervals, the lungs were harvested 3 days after RSV challenge. Comparison of Adju-Phos-containing KD-409 and adjuvant-free pre-F; (B) anti-F antibody titer; (C) neutralizing antibody titer; and (D) protective efficacy against infection. Statistical analysis was performed by one-way-ANOVA and Dunn’s multiple comparison test. n = 9-10; **p < 0.01, *p < 0.05, ns: p > 0.05, not significant.

To confirm dose-dependent immunogenicity and efficacy, we measured the anti-F and neutralizing antibody titers in blood and viral copy numbers in lung tissue after RSV challenge. Immunization with a 0.5–5.0 µg/dose of KD-409 induced dose-dependent anti-F antibody titers; the adjuvanted antigen amount of 1/10 of KD-409 had a higher anti-F antibody induction ability than the adjuvant-free pre-F (Fig. 4B). In terms of neutralizing antibody titers, KD-409 with adjuvant significantly outperformed pre-F without adjuvant, despite a 1/10 antigen dose (Fig. 4C). Assessing protection against infection in the cotton rat model, KD-409 with an adjuvant significantly outperformed pre-F without an adjuvant, which was five times the amount of antigen. Adjuvanted KD-409 reduced viral copy numbers (GMT) compared to the unadjuvanted pre-F tenfold antigen dose (Fig. 4E). These results suggest that KD-409 is superior to adjuvant-free pre-F in terms of efficacy in the cotton rat model.

## DISCUSSION

In this study, we evaluated the safety and efficacy of KD-409, a chimeric RSV FG protein that incorporates a highly conserved region of the RSV G protein, in various rodent species. In a passive immunization model in which pregnant females were inoculated, we found that antibody transfer occurs in guinea pig models and that KD-409 is less likely to exacerbate infection in a mouse passive model than conventional antigens such as FI-RSV and pre-F with Adju-Phos. In a low-dose mouse model of active immunization that mimics infant vaccination and fails to induce sufficient immunity, KD-409 with Adju-Phos was less likely to exacerbate inflammation than FI-RSV or pre-F with Adju-Phos. In a cotton rat model, which is a well-established method for evaluating the efficacy of human RSV vaccines, KD-409 was significantly more effective in eliciting protection against infection than unadjuvanted pre-F.

Currently, there is no RSV vaccine covering from neonates to children, that is both effective and safe. When pregnant women are vaccinated, transferable antibodies from the mother to the neonate are expected to decline over time. Regardless of vaccination, the neutralizing antibody titer at 3 months of age is less than 1/10 of that at 0 days of age (22), and 60% of hospitalizations due to RSV infection in children under 1 year of age occur within the first 3 months of life (23). Moreover, vaccination of children may not induce sufficient immunity because of the characteristics of childhood immunity (24). Moreover, vaccination of pregnant women alone is not sufficient to prevent severe illness and hospitalization in newborns, infants, and children. RSV vaccines need to be developed for both newborns with decreasing maternal antibodies and children with immature immune systems. Given that VED occurred approximately half a century ago (2), vaccine development for RSV in children has progressed with caution, and there are no licensed vaccines. Therefore, vaccine efficacy and safety concerns exist and need to be addressed for both maternal and pediatric vaccines.

We previously demonstrated the immunogenicity and efficacy of KD-409 in mouse models of active and passive immunization (18). In the current study, we focused on the safety of KD-409 for maternal and pediatric immunization and compared its efficacy to that of the approved formulation, pre-F, without an adjuvant for maternal immunization. In the case of tetanus and pertussis vaccines administered to pregnant women, the half-live of transferable antibodies is approximately 30 days for both (17). The half-life of the maternal RSV antibody is 36–42 days (25), and the neutralizing titer of the transfer antibody induced by RSVPreF3 maternal immunization decreases by half within 43 days (22). In the guinea pig models, anti-F antibody and neutralizing antibody titers were halved within 30 days of birth with the alum-adjuvanted RSV recombinant F nanoparticle vaccine (20), consistent with our findings (Fig. 1H). The half-life of transplacental antibodies induced by KD-409 is expected to be similar to that of conventional maternal immunization. When antibodies are transferred from mother to newborns, the concentration of antibodies increases, a process known as antibody concentration (26). In our experiment, the anti-F antibody titer (GMT) of the pups was higher than that of the mother on day 0. This result suggests that the concentration of transitional antibodies induced by KD-409 is similar to that induced by conventional maternal immunization.

Cases of severe RSV infection in infants requiring hospitalization peak at approximately 2–3 months of age, and the risk of severe RSV infection is also high in those whose transient antibodies have lost their effectiveness and in children under 5 years of age (1, 15, 16, 27, 28); hence, an age-appropriate vaccine is urgently needed. Children have immature immune systems, and vaccination may fail to induce sufficient antibody titers (29). *Ex vivo* and *in vitro* studies have suggested that inadequate immunity may result in low neutralizing antibody titers, increasing the likelihood of enhanced ADE infection (30). In an active immunization model of cotton rats, low-dose immunization with the F antigen alone demonstrated a trend toward the exacerbation of symptoms after vaccination with post-F or pre-F with alum (31). In addition, immunization with pre-F-alum exacerbated symptoms in a BALB/c mouse model (9). A new and simplified *in vivo* system is required to evaluate infection-induced adverse reactions and symptom exacerbation after vaccination. In this study, we demonstrated that KD-409 with Adju-Phos is unlikely to cause infection and symptom exacerbation after vaccination using a mouse passive immunization model and a low-dose mouse active immunization model (Figs. 2 and 3). Lung sections from infants who died in the 1967 FI-RSV clinical trial identified respiratory eosinophils and CCL5 as markers of enhanced RSV disease (32). In this study, we focused on eosinophils and evaluated them in a mouse model, demonstrating that pre-F-Adju-Phos increased eosinophil counts, whereas KD-409-Adju-Phos did not. In this study, we demonstrated that KD-409 was less likely to exacerbate infections and symptoms (Figs. 2 and 3). We also discuss the mechanism of exacerbation suppression. CX3C in RSV G suppresses the function of normal Th1 cells by interacting with CX3CR1 in the host cell, whereas anti-G antibody promotes Th1 cell function by inhibiting this interaction (33). In a cell migration assay, 3D3 antibodies functionally inhibited the CX3C-CX3CR1 interaction (34). As reported previously, KD-409 inoculation results in Th1 dominance; however, the induced anti-G antibody titer remains relatively low (18), making it difficult to verify in the cell migration system. Whether low anti-G antibody levels are a significant factor in the inhibition of exacerbation remains debatable. We believe that the various immune responses to KD-409 need to be examined; hence, we are actively acquiring additional data.

The occurrence of infection and symptom exacerbation may depend on the antibody concentration in the blood; however, it is difficult to measure all antibody concentrations comprehensively, which is a limitation of this experimental system. ABRYSVO, approved for vaccination of pregnant women, is an adjuvant-free pre-F. This approved vaccine contains bivalent pre-F antigens against RSV A and B strains mutated at 847aa to enhance conformational stability and immunogenicity (35). This antigen differs from previously reported DS-Cav1 mutants, including S155C, S290C, S190F, and V207L (36). It is important to note that the pre-F used in our experiments is a monovalent DS-Cav1 mutant vaccine derived from the RSV A strain; therefore, our results are not strictly comparable to those of the licensed vaccine (Fig. 4). Given that the Th2-primed immune system is considered to be predominant in neonates and children (37), and that BALB/c mice are Th2-predominant compared to C57BL/B6 mice (38), we used this animal model to mimic pediatric vaccination in this experiment. However, the biological indicators, except for Th2 dominance, differ from those of human newborns and children. Moreover, although the cotton rat model is an established RSV vaccine evaluation system, from the perspective of human extrapolation, a monkey model is more desirable for evaluating the safety and efficacy (39). To verify the high accuracy of safety and efficacy, a test using the monkey model will probably be necessary in the future, taking into account the extrapolation to humans.

In conclusion, we evaluated the next-generation RSV vaccine KD-409 in rodent models and demonstrated that its safety and efficacy were superior to those of the pre-F antigen. Accordingly, KD-409 may be suitable for both maternal and pediatric vaccination. KD-409 is a promising next-generation RSV vaccine for neonates and children.

## MATERIALS AND METHODS

### Antigens, viruses, and cells

FI-RSV, pre-F, KD-409, and RSV A2 (VR-1540; ATCC, Manassas, VA, USA) were prepared as reported previously (18). FI-RSV and RSV A2 were obtained from Hep-2 (CCL-23; ATCC); pre-F was obtained from Expi293 (A14527, Thermo Fisher Scientific K.K., Tokyo, Japan), and KD-409 was obtained from Expi293 or CHO DG44 (Meiji Seika Pharma). Further details are available in our previous report (18).

### Immunization

#### Mouse (active)

Six- to seven-week-old female BALB/c mice (Japan SLC, Inc., Shizuoka, Japan) were administered two intramuscular doses of various antigens at 3-week intervals. Three weeks later, blood was collected under isoflurane anesthesia and centrifuged (3000×g, 10 min, 25°C) to obtain serum.

#### Mouse (passive)

The sera of other actively immunized mice were diluted with phosphate-buffered saline and administered intraperitoneally.

#### Guinea pig (pregnant)

Pregnant guinea pigs (Japan SLC) were administered two intramuscular injections of antigens at 3-week intervals, and their blood, together with that of the pups, was collected 0, 15, and 30 days after delivery. The collected blood specimens were centrifuged (3000×g, 10 min, 25°C) to obtain serum.

#### Cotton rats (active)

Female Hsd cotton rats (ENVIGO/Japan SLC) aged 11–12 weeks were administered two intramuscular injections of antigens at 3-week intervals. The collected blood specimens were centrifuged (3000×g, 10 min, 25°C) to obtain serum.

### Enzyme-linked immunosorbent assay (ELISA) and cell-ELISA

#### Anti-F antibody titer

Serum was applied to 96-well Pierce nickel-coated plates (Thermo Fisher Scientific K.K., Tokyo, Japan), immobilized with pre-F, and blocked with 1% bovine serum albumin. Detection was performed using horseradish peroxidase-conjugated anti-mouse IgG (Global Life Sciences Technologies Japan K.K., Tokyo, Japan), anti-guinea pig IgG, anti-cotton rat IgG, and 3,3′,5,5′-tetramethylbenzidine liquid substrate (Sigma-Aldrich Co., St. Louis, MO, USA). Further details are available in our previous report (18).

#### Neutralizing antibody titers

In brief, 2 × 10^5^ cells/mL HEp-2 cells (CCL-23; ATCC) were pre-cultured in 96-well plates. Serum and RSV diluent containing rabbit serum complement (Cedarlane, Ontario, Canada) were mixed in equal volumes and incubated at 37℃ and 5% CO_2_ for 1 h. After removing the culture supernatant from the plate, the serum-RSV reaction solution was added and incubated at 37℃ and 5% CO_2_ for 2 days. The infected cells were fixed in methanol, air-dried, and detected with an anti-F antibody, anti-mouse IgG-Alexa488 conjugated antibody (Abcam, Cambridge, UK), and Hoechst 33342 solution (Dojindo Laboratories Co., Ltd., Kumamoto, Japan). The infection rate was analyzed using an image analyzer (Image Xpress Micro XLS; Molecular Devices, LLC., San Jose, CA, USA), and the neutralizing antibody titer (IC_50_) was calculated by curve fitting in GraphPad Prism 9.5.1 (GraphPad Software Inc., San Diego, CA, USA) or set by the dilution factor that exceeded a 50% inhibition rate. Further details are available in our previous report (18).

### Virus challenge and copy number analysis

RSV A2 (1 × 10^5^–10^6^ PFU) was inoculated intranasally (10–20 µL or 100 µL; mouse-active/pregnancy or cotton rat) under 2%–3% isoflurane anesthesia. Three days after viral challenge, whole lungs were collected in Lysing Matrix D tubes (MP-Bio Japan K. K., Tokyo, Japan), followed by the addition of TRIzol (Thermo Fisher Scientific K.K.). The tubes were agitated with FastPrep-24 (MP-Bio Japan K.K.), and the lung tissue was crushed. cDNA was synthesized from the extracted RNA using a High-Capacity cDNA Reverse Transcription Kit (Thermo Fisher Scientific). Quantitative polymerase chain reaction (PCR) was performed to detect the number of viral copies using the sense primer CARCAAAGTTAYTCTATCATGTC, antisense primer GATCCTGCATTRTCACARTACCA, minor groove binder (MGB) probe TGTAGTACAATTRCCACT, and standard DNA; TGTCCAACAATGTTCAAATAGTTAGACAGCAAAGTTACTCTATCATGTCCATAATAAAAGAGGAAGTCTTAGCATATGTAGTACAATTACCACTATATGGTGTTATAGATACACCCTGTTGGAAACTACACACATCCCCTCTATGTACAACCAACACAAAAGAAGGGTCCAACATCTGTTTAACAAGAACTGACAGAGGATGGTACTGTGACAATGCAGGATCAGTATCTTTCTTCCCACAAGCTGAAACA TGTA. Further details are available in our previous report (18).

### Inflammation assessment

#### Lung tissue section

BALB/c mice were immunized twice with antigen protein containing 0.5, 5, 50, or 500 ng/dose with Adju-Phos and challenged with RSV A2. Whole lungs were harvested 3 days after the challenge, and the left lung was fixed in 10% formalin. Paraffin-embedded tissues were sectioned at a thickness of 2.5 µm and stained with hematoxylin and eosin for histological evaluations. A total of four mice were assigned to each dose (n = 4), and three tissue sections were prepared for each mouse. The results of the group exhibiting the highest inflammation level were used. Pulmonary inflammatory symptoms (perivascular interstitial edema, peribronchial and perivascular cellular infiltration, alveolar wall thickening, and bronchial wall mucus) were evaluated by an experienced specialist using an Eclipse 80i microscope (NIKON CORPORATION, Tokyo, Japan).

#### Inflammatory cells

BALB/c mice were immunized twice with an antigen protein containing 5 ng/dose with Adju-Phos were challenged with RSV A2. The BALF was collected 3 days after the challenge, and cells were applied to glass slides using Cytospin 4 (PHC Corporation, Tokyo, Japan). Slides were stained with Diff-Quik (16920; SYSMEX CORPORATION, Kobe, Japan). The numbers of multinucleated cells, eosinophils, and neutrophils were counted by a trained specialist using ImageXpress pico (Molecular Devices, LLC., San Jose, CA, USA).

### Statistical analysis

Anti-RSV F antibodies, neutralizing antibody titers, and viral loads are presented as geometric means and 95% confidence intervals (CIs). The ratios of PMN leukocytes, eosinophils, and neutrophils are presented as arithmetic means and 95% CIs. Statistical analysis was performed using GraphPad Prism version 9.5.1 (GraphPad Software Inc.). Neutralizing antibody titers and virus copy numbers were analyzed using the Kruskal–Wallis test and Dunn’s multiple comparison test. For the evaluation of inflammation, the Brown-Forsythe test, Welch’s ANOVA, and Dunnett’s T3 multiple comparison test were used.

## Data availability Statement

The datasets presented in this article are not readily available because the data are part of an ongoing study. Requests to access the datasets should be directed to the corresponding author.

## Acknowledgments

We are grateful to Meiji Seika Pharma Co. Ltd., Hiroaki Mori, Hiromi Nakamura, Sayaka Mizumoto, Keiko Goto, Minako Inui, Mika Kiyokawa, Yuki Koga, Satomi Harano, Noriko Shinya, Tomohiro Nishimura, and Hirofumi Higuchi for their assistance with this study.

## Author contributions

Conceptualization, R.Y., M.T. (Madoka Terashima), M.T. (Masaharu Torikai), and K. S.; formal analysis, R.Y., M.T. (Madoka Terashima), and M.T. (Masaharu Torikai); investigation, R.Y. and M.T. (Madoka Terashima); writing-original draft preparation, R.Y. and M.T. (Masaharu Torikai); writing-review and editing, R.Y., M.T. (Madoka Terashima), M.T. (Masaharu Torikai), and K. S. All authors have read and agreed to the published version of the manuscript.

## Institutional review board statement

The animal study protocol was approved by the Institutional Animal Care and Use Committee of KM Biologics.

